# Divergent Effects of Ultra-High Dose Arachidonic Acid versus Docosahexaenoic Acid on *MYCN*-Driven Neuroblastoma Progression in a Syngeneic Mouse Model

**DOI:** 10.1101/2025.05.06.652504

**Authors:** V Patel, YN Li, X Shen, JT Brenna, JT Powers

## Abstract

Neuroblastoma (NB) represents the most common extracranial solid tumor in children, where high-risk cases have particularly poor prognosis. We used a syngeneic mouse model to investigate the effects of ultra-high dose highly unsaturated fatty acids (HUFA) on NB progression. Following tumor establishment, mice were randomized to receive daily oral gavage with omega-6 (ω6) arachidonic acid (ARA) at 4.7 g/d human equivalent (hEq), or omega-3 (ω3) docosahexaenoic acid (DHA) at 24 g/d hEq, and controls did not receive gavage. We observed strikingly divergent effects: ARA significantly promoted tumor growth, resulting in 100% tumor survival and 4-fold larger tumors compared to controls, with enhanced vascularization and invasive morphology. In contrast, DHA administration reduced tumor survival (40% versus 92% in controls) and significantly suppressed the progression of remaining tumors, with remaining DHA-treated tumors approximately 4.5-fold smaller than controls and 18-fold smaller than ARA-treated tumors. In a separate lipid mediator analysis, ARA supplementation significantly increased pro-inflammatory/pro-tumorigenic mediators including PGE2, TXB2, and epoxyeicosatrienoic acids in liver, spleen, brain and skeletal muscle. DHA supplementation increased anti-inflammatory/anti-tumor mediators, particularly EPA-derived 17,18-EpETE, 18-HEPE, and DHA-derived 14-HDHA in these same tissues. No significant differences in body weight were observed among treatment groups, indicating the treatments were well-tolerated. These findings build upon our previous research demonstrating that ultra-high dose ω3 supplementation can completely block tumor formation in MYCN-driven NB. The profound tumor-suppressive effects of DHA suggest that dietary modulation of ω3 and ω6 HUFA intake may offer a complementary, low-toxicity approach to high-risk NB standard of care (SoC). Our findings suggest that dietary intervention may be an effective primary or adjunctive strategy in pediatric oncology, enabling reduced SoC dosing while improving outcomes and survivorship.

## Introduction

Neuroblastoma is an aggressive pediatric cancer originating from neural crest-derived progenitor cells, representing the most common extracranial solid tumor in children with a disproportionate mortality rate(1). Clinically, it exhibits remarkable heterogeneity, from spontaneously regressing tumors to rapidly progressing, treatment-resistant forms. Inflammation has emerged as a critical component in tumor biology, playing key roles in initiation, progression, and immune evasion (2).

The tumor microenvironment, characterized by its inflammatory mediators, cytokines, and bioactive lipids, significantly influences tumor behavior. Polyunsaturated fatty acids (PUFAs), particularly those belonging to the omega-6 and omega-3 families, play opposing roles in cancer biology.(3) Arachidonic acid (ARA), a primary omega-6 PUFA in animal-derived phospholipids, generates pro-inflammatory eicosanoids that promote angiogenesis, suppress anti-tumor immunity, and enhance metastasis.(4) Higher levels of ARA metabolites correlate with aggressive tumor phenotypes and poorer outcomes across multiple cancers, including neuroblastoma. In contrast, omega-3 fatty acids, primarily eicosapentaenoic acid (EPA) and docosahexaenoic acid (DHA), exhibit anti-inflammatory properties and offer numerous health benefits. These essential fatty acids compete with ARA in cell membrane incorporation and enzymatic pathways, reducing pro-inflammatory eicosanoids and possibly increasing anti-inflammatory mediators like resolvins and protectins (5). Clinical applications have shown that ultra-high dose omega-3 therapy (up to 20 g/day) is well-tolerated and effective in conditions including traumatic brain injury and hypertriglyceridemia (6).

Preclinical evidence suggests that omega-3 fatty acids inhibit tumor cell proliferation, induce apoptosis, and modulate immune responses (7–9). Notably, our recent publication demonstrated that ultra-high doses of oral omega-3 fatty acids (EPA, DHA, or oxidation-resistant deuterated DHA) at human-equivalent doses of 12-14 g/day completely blocked tumorigenesis in a MYCN-driven neuroblastoma model(10). Conversely, omega-6 ARA supplementation enhanced tumor formation and reduced tumor latency, highlighting the opposing impacts of these lipid families (11)

We report here an investigation of the therapeutic potential of DHA in a syngeneic mouse model of MYCN-driven neuroblastoma, examining its effects on tumor growth kinetics to provide insights for potential dietary interventions in pediatric cancer management.

## Methods

### Cell Culture and MYCN Transfection

Murine neuroblastoma Neuro-2a cells transduced with a human *MYCN* transgene were initially obtained from the American Type Culture Collection (ATCC; CCL-131; details available at ATCC) and cultured in RPMI-1640 medium supplemented with 10% heat-inactivated fetal bovine serum (HI-FBS). Cells were grown at 37 °C in a water-saturated atmosphere of 95% air and 5% CO_2_. Cells were tested monthly and consistently found to be negative for mycoplasma contamination. Cells were stably transfected with human MYCN using Lipofectamine 3000. Transfected cells were selected from GFP-expressing cells using a fluorescence-activated cell sorter.

### Animal Model and Treatment

All procedures were IACUC-approved. Six-week-old male wild-type strain A/J mice (n = 42) were purchased from The Jackson Laboratory (Jaxmice strain #000646, Sacramento, CA, USA) and allowed to acclimate for one week under standard conditions. Mice were fed customary mouse chow with a composition approximating AIN-93G. A model was established by subcutaneous injection of 3 × 10^6^ MYCN-expressing Neuro-2a cells into the right flank of each mouse. When tumors became palpable (approximately day 6), mice were randomized into three treatment groups: Control group (n=12), ARA group (n=15) receiving daily oral gavage with 75 µL ARA oil (44% ARA, 4.6 g/d human equivalent (hEq) dose), and DHA group (n=15) receiving daily oral gavage with 250 µL DHA oil (82% DHA, 24 g/d hEq). All dose and chow fatty acids were as presented previously (10). Oral gavage was performed daily throughout the study. Tumor dimensions were measured every two days using digital calipers, with volume calculated as: Tumor volume = (length × width^2^) × 0.5. Body weight and health status were continuously monitored. The study concluded when the control tumors reached approximately 1,000 mm^3^. Mice were euthanized by CO_2_ inhalation followed by cervical dislocation. Tumors were excised, photographed, weighed, and processed for analysis (flash-frozen, fixed in formalin, or processed immediately).

#### Lipid mediator experiment

We performed a separate experiment to understand the effects of ARA and DHA dosing on eicosanoid and docosanoid production in key tissues: liver, spleen, brain, and skeletal muscle. Mice were gavaged daily with the same levels of DHA and ARA, or no gavage, with n=7 per group. At 7 days, 1 × 10^6^ MYCN-expressing Neuro-2a cells were injected into the bloodstream as an alternative model. No tumors developed in these animals as assessed by inspection of the liver and lung and by mouse behavior. Mice were gavaged daily until 62 days of life when they were euthanized. Liver, spleen, brain and skeletal muscle tissues are harvested from 12 male and 9 female mice, and dropped immediately into 1 ml of ice-cold methanol containing 10ng/ml of spiked internal standard. Samples are homogenized using a ground glass pestle tissue grinder until fully homogenized or until the solvent appears cloudy. The tube containing the tissue and solvent is vortexed and subsequently incubated on ice for 10 minutes. 9mls of water is added to dilute methanol composition to 10%. Strata-X 33 µm Polymeric Reversed Phase cartridges purchased from Phenomenex (California, USA) are used for solid phase extraction (SPE), the final eluded samples are dried under a gentle stream of nitrogen, and reconstituted with 60ul of 50/ 50 MeOH/ Water (v/v), before being placed in the autosampler.

### Lipid mediators

Lipid mediator detection was performed using a 20-minute scheduled multiple reaction monitoring (sMRM) method on a Kinetex 2.6 µm Polar C18 100 Å column (100 mm × 3 mm) coupled to a Sciex 7500 triple quadrupole QTRAP mass spectrometer operating in negative electrospray ionization mode, integrated with an ExionLC-AD UHPLC system. Chromatographic separation was performed on a Kinetex 2.6 µm Polar C18 100 Å column (100 mm × 3 mm) with column oven temperature at 50°C. Collision energy (CE) and exit potential (CXP) were optimized through direct infusion using available standards. Electrospray ion source and gas parameters were optimized through on-column flow injection analysis (FIA). The LC flow rate is stable at 0.5ml/min with mobile phase A consisting of 100% water, and mobile phase B consisting of ACN/ MeOH 60/40 (v/v), each containing 0.5% of HAc as modifier. The LC gradient is as follows: 0-2mins at 45% B, 2-5mins increase to 48% /B, 5-16.5mins increase to 80%, 16.6min increase to 98%, hold at 98% until 18.6mins then return to 10% B until 20.5min. 74 lipid mediators were included in the final sMRM method, target cycle time is at 700ms, minimum and maximum dwell time at 3 and 500ms, respectively.

Analytical standards were Primary Vascular Eicosanoid MaxSpec® LC-MS Mixture (Cayman Chemicals No.19667), Arachidonic Acid CYP450 Metabolite MaxSpec® LC-MS Mixture (Cayman Chemicals No.20665), Primary COX and LOX MaxSpec® LC-MS Mixture (Cayman Chemicals No.19101), Arachidonic Acid Oxylipin MaxSpec® LC-MS Mixture (Cayman Chemicals No.20666), SPM E-series MaxSpec® LC-MS Mixture (Chemicals No.19417). Prostaglandin E1 (Cayman Chemicals No.13010), Prostaglandin E3 (Chemicals No.14990), Leukotriene B4 (Cayman Chemicals No.20110), DHA Oxylipin MaxSpec® LC-MS Mixture (Cayman Chemicals No.22280), SPM D-series LC-MS Mixture (Cayman Chemicals No.18702), SPM E-series LC-MS Mixture (Cayman Chemicals No.19417), 8-iso Prostaglandin F2a (Cayman Chemicals No. 16350), Lipoxin LC-MS Mixture (Cayman Chemicals No. 19412), and Docosahexaenoic Acid CYP450 Oxylipins LC-MS Mixture (Cayman Chemicals No. 22639).

#### Data analysis

For each sample, chromatographic peaks were integrated to obtain the area under the curve (AUC) for each target analyte and for the 5-HETE-d8 internal standard (IS). Peak integration is conducted through the auto-peak algorithm in the Analytics software from Sciex-OS, each integration is then checked manually. An analyte to IS ratio is calculated by dividing the analyte’s AUC by the IS’s AUC, to compensate for instrument performance differences between injections. The median of the analyte/ IS values within a sample is calculated for all samples. A median normalized ratio is calculated for every analyte by dividing its analyte/IS ratio by the sample’s median value. This normalization sets the median analyte/IS ratio within each sample to be 1, centering the data and allows comparability across samples. Next, the mean of the median normalized ratios in the control samples are calculated for each analyte to provide a baseline for comparison between treatments. Fold changes relative to the control samples are determined by their median-normalized ratio by the corresponding control-group mean. Thus, a treatment sample with a fold-change of 1.0 indicates an analyte level equivalent to the control mean, whereas a fold-change of 2.0 or 0.5 would indicate a level double or half that of the control mean, respectively. Lipid mediators are reported as mean ± SD. Statistical significance at p<0.05 was determined with Welch’s t test with no correction for multiple comparisons.

### Statistical Analysis

Data are presented as mean ± SEM. Analyses were performed using GraphPad Prism (version 3.1). Comparisons between groups used one-way ANOVA with Tukey’s post hoc test. For tumor growth kinetics, two-way repeated measures ANOVA were used. P-values < 0.05 were considered significant.

## Results

### DHA suppresses tumor growth in a MYCN-driven neuroblastoma model

To evaluate the therapeutic potential of omega-3 fatty acid (DHA) in neuroblastoma, we employed a syngeneic xenograft model using MYCN-transfected Neuro2a cells. Mice (n=42) were injected subcutaneously with Neuro-2a cells. At 6 days, the 30 mice with palpable tumors were randomly assigned to one of three groups so that each group had mice with 10 tumors. The remaining mice were assigned randomly to groups to make the initial treatment groups as follows (***Figure 1***)—Control (n=12), ARA (omega-6, n=15), and DHA (omega-3, n=15). At day 8, more mice had developed palpable tumors to make the final count of tumors per group as Control (11/15), ARA (11/15), DHA (15/15) when daily oral gavage started on day 8 post-injection. One cage containing several mice were euthanized due to wounds acquired from fighting over the course of 29 d of gavage (Control, 1; DHA, 3; ARA, 3), leaving at study termination the following numbers of mice that ever had tumors: Control 10, ARA 12; DHA 12. At the day 37 study termination, tumor counts were control, 10/10, ARA 12/12, DHA 6/12; in the DHA group, 6 mice that had tumors were tumor-free by palpation.

**Figure 1.**
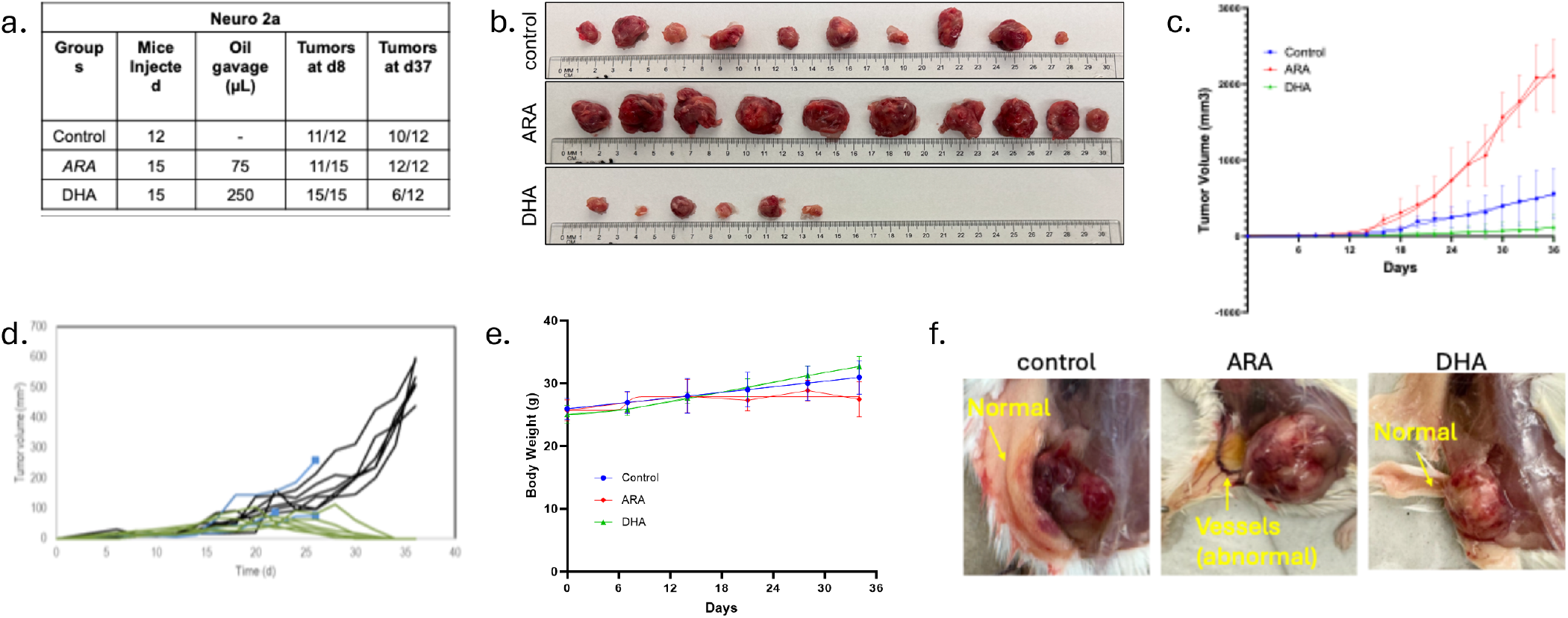
Ultra-high-dose DHA inhibits tumor growth in a MYCN-driven neuroblastoma model. **a)** Table representing the results of subcutaneous injection of 3×10^6^ *MYCN*-transfected Neuro-2a cells followed by treatment administered via oral gavage daily, starting 8 days post-injection. b) Representative images of tumors harvested from each group at the end of the study (day 36). Tumors were aligned along a metric scale to illustrate size differences between treatment groups. **c)** Tumor growth curves over time. Tumor volume (mm^3^) was measured every 2 days starting from day 8. Data represent mean ± SEM; n = 12–15 mice per group. **d)** Individual DHA-treated tumor growth curves over time. **e)** Body weight of mice during treatment period. No significant differences in body weight were observed among the groups, indicating minimal. **f)** Gross morphology of subcutaneous tumors at endpoint highlighting differences in vascularization and surrounding tissue architecture.

At the endpoint (day 37), tumors were collected and aligned for visual comparison (***Figure 1b***). The DHA group displayed drastically smaller and fewer tumors, whereas the ARA group exhibited the most significant tumor burden. Longitudinal monitoring of tumor progression (***Figure 1c***) revealed that DHA treatment nearly abolished tumor growth, while ARA significantly accelerated tumor progression compared to controls. Mice receiving ARA exhibited dramatically accelerated tumor growth compared to the control group, with tumor volumes approximately 4-fold higher by Day 37 (p<0.01). In contrast, DHA treatment substantially inhibited tumor progression, with mean tumor volumes approximately 4.5 times smaller than those of the control group (p < 0.01) and 18 times smaller than those of the ARA group (p < 0.001) by the endpoint. Individual tumor growth was broadly suppressed by DHA, including six tumors that regressed completely (***Figure 2d***).

**Figure 2.**
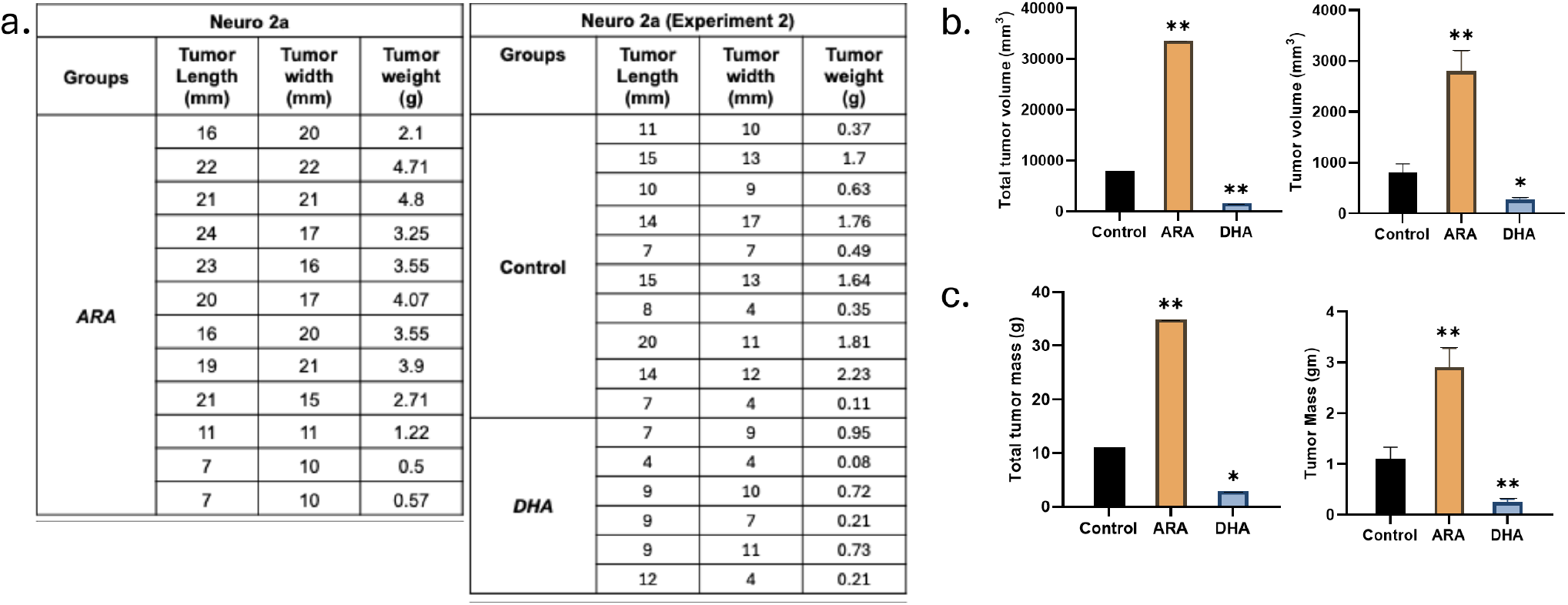
Syngeneic cell-derived xenograft model of *MYCN*-driven neuroblastoma. ARA and DHA were administered orally every day to the mice after 8 days (tumor palpable days) after cell injection. 3 ^*^10^6 *MYCN expressing* Neuro-2a cells were injected subcutaneously into the mice**—summary** of tumor size and mass with ARA, DHA, and control. A bar graph depicts the change in mean tumor mass across different experimental groups, showing the highest tumor volume in the ARA-treated group in both experiment rounds. Significance was calculated using a t-test: ^*^p<0.05 and ^**^p<0.01.

Importantly, no significant differences in body weight were observed among the treatment groups (***Figure 1e***), indicating that the treatments were well tolerated, with some evidence of divergence at the end of the study. The gross site of tumors was inspected (***Figure 1f***), revealing that ARA-treated tumors appeared more vascularized and invasive, with irregular borders and evidence of abnormal vessel formation. In contrast, tumors from DHA-treated mice were generally well-circumscribed, paler in appearance, and showed standard surrounding tissue architecture. Control tumors exhibited intermediate characteristics. DHA reduces tumor volume and mass across experimental replicates.

Detailed analysis of tumor dimensions and mass (***Figure 2a***) further confirmed the differential effects of ARA and DHA treatments. This analysis included comprehensive measurements of individual tumors from each treatment group, revealing consistent patterns. The mean tumor length and width in the ARA group (20.00 mm × 17.83 mm) significantly exceeded those in both the control (16.00 mm × 20.00 mm) and DHA groups (7.00 mm × 10.00 mm). Quantitative analysis of the tumor metrics revealed that ARA treatment resulted in the highest total tumor volume and mass, whereas DHA treatment yielded a significant reduction in both parameters. Bar graphs summarizing the tumor volume (***Figure 2b***) and tumor mass (***Figure 2c***) confirmed that DHA significantly suppressed tumor growth (*p* < 0.05 vs. control, *p* < 0.01 vs. ARA). At the same time, ARA treatment resulted in significantly larger tumors (*p* < 0.01 vs. control). The total tumor mass in the ARA group was approximately three times higher than in the controls and six times higher than in the DHA group.

Together, these data demonstrate that DHA causes regression or markedly reduces tumor progression in *MYCN*- driven neuroblastoma, while ARA promotes tumor growth and abnormal tissue morphology. Importantly, DHA achieved these anti-tumor effects without causing systemic toxicity, suggesting its potential as a safe therapeutic approach for neuroblastoma treatment.

### Lipid Mediators

Eicosanoid and docosanoid levels for the ARA- and DHA-dosed animals are reported as fold-changes compared to the Control. We report summary data consisting of changes in lipid mediators that were significantly different between ARA- and DHA-dosed animals (Tables 1-3). All EPA and ARA-derived eicosanoids showed significant changes between ARA and DHA dosing in at least one tissue, while for DHA-derived docosanoids 12 of 20 were significantly changed. In all cases, the direction of changes favored the dosed fatty acid, except for LTB4 in liver. In DHA dosed animals, all 10 EPA-derived eicosanoids changes in the expected direction in liver, and 9, 8, and 7 EPA-derived eicosanoids changed in spleen, brain, and skeletal muscle, respectively.

**Table 1.** EPA-derived eicosanoids in tissue. Ratio of DHA-dosed to ARA-dosed oxylipins for those that are p<0.05. Comparison of ARA and DHA dosed animal fold-changes compared to controls as described in the Methods. MRM transitions are reported as (precursor/product) masses. – means not significant

| EPA Derived Eicosanoids | Liver | Spleen | Brain | Skeletal Muscle |
| --- | --- | --- | --- | --- |
| 8,9-EpETE (317/127) | 4.69 | 3.21 | 2.19 | - |
| 11,12-EpETE (317/167) | 3.90 | 3.90 | 3.36 | 1.85 |
| 14,15-EpETE (317/207) | 4.60 | 3.54 | - | 1.94 |
| 17,18-EpETE (317/215) | 4.93 | 2.62 | 2.71 | 2.14 |
| 17,18 DiHETE (335/203) | 2.08 | - | - | 1.85 |
| 18-HEPE (317/259) | 4.87 | 3.02 | 2.84 | 1.87 |
| 5-HEPE (317/115) | 4.04 | 2.31 | 2.27 | 2.58 |
| 11-HEPE (317/167) | 3.62 | 2.85 | 2.65 | - |
| 12-HEPE (317/179) | 2.68 | 3.49 | 2.60 | 2.54 |
| 15-HEPE (317/219) | 2.68 | 1.88 | 2.14 | - |

ARA-derived PGE2, PGD2, and PGF2a all changed in liver, PGF2a and the isoprostane 8-iso-PGF2a changed in skeletal muscle. Changes in ARA-derived eicosanoids in the Brain were limited to the 8,9-diHEtrE and both diHETs. In

## Discussion

Neuroblastoma, particularly in high-risk *MYCN*-amplified cases, remains a therapeutic challenge due to its aggressive progression and treatment resistance. Our study reveals striking divergent effects of ultra-high omega-6 and omega-3 fatty acids on neuroblastoma progression: ARA significantly promoted tumor growth, while DHA consistently suppressed it. The tumor-promoting effect of ARA aligns with its role as a precursor to pro-inflammatory eicosanoids that enhance angiogenesis, immune evasion, and metastasis (12). The cyclooxygenase-2 (COX-2) pathway, which metabolizes ARA to prostaglandins, is frequently upregulated in various cancers, including neuroblastoma, with pathway inhibition showing anti-tumor effects (13).

Our data provides compelling in vivo evidence of ARA’s tumorigenic activity in neuroblastoma. The dramatic tumor growth enhancement observed with daily ARA administration (75 µL/day, 44% ARA) resulted in 4-fold larger tumors compared to controls and 100% tumor incidence. These results suggest that even modest increases in dietary omega-6 intake may significantly accelerate disease progression. Notably, ARA-treated tumors displayed abnormal vascularization and invasive morphology, consistent with enhanced angiogenesis and metastatic potential. This finding is corroborated by our previous study, which demonstrated that ARA supplementation (4.6-6.0 g/day, human dose equivalent) enhanced tumor formation, increased vascularization, and reduced tumor latency from 10 to 5.5 days in a similar model (6). These results have important implications for dietary recommendations in pediatric cancer patients, especially given the high omega-6 content in Western diets. Conversely, DHA (250 µL/day, 100% DHA) exerted a powerful tumor-suppressive effect, significantly reducing both tumor incidence (40% in DHA vs. 92% in controls) and tumor progression. Tumors that did develop in DHA-treated mice were approximately 4.5-fold smaller than controls and 18-fold smaller than ARA-treated tumors. Notably, DHA treatment was associated with well-circumscribed tumors characterized by standard surrounding tissue architecture, suggesting effects extending beyond simple growth inhibition. These findings align with multiple preclinical studies reporting anti-proliferative, pro-apoptotic, and immune-modulating effects of omega-3 fatty acids.

Also, several mechanisms may contribute to DHA’s anti-tumor effects: (1) competitive inhibition of ARA metabolism, reducing pro-inflammatory eicosanoid production; (2) alteration of lipid raft composition; (3) direct modulation of gene expression through nuclear receptors; and (4) potential production of specialized pro-resolving mediators that actively resolve inflammation. (8,9,14,15) The opposing effects of ARA and DHA are particularly relevant in *MYCN*-driven neuroblastoma, where *MYCN* amplification is associated with poor prognosis and accelerated tumor growth. *MYCN* reprograms cellular metabolism, including lipid metabolism, to support rapid proliferation (16). Our findings suggest that dietary fatty acids interact with MYCN-driven metabolic alterations, either exacerbating (ARA) or mitigating (DHA) their effects on tumor progression.

### Effects of Lipid Mediators on MYCN and MYC-Associated Cancers

Our MYCN-amplified Neuro-2a cell-induced tumors most closely model aggressive MYCN-amplified neuroblastoma. We first consider a subset of known effects of the lipid mediators whose concentrations changed between ARA and DHA dosing in MYCN-amplified disease, then consider MYCN-non-amplified and MYCN-amplified disease other than neuroblastoma. The known functional redundancy of MYCN and MYC implies similarity of effects in the wide range of cancers associated with MYC-associated cancers.

### MYCN-Amplified Neuroblastoma

MYCN-amplified neuroblastoma represents one of the most aggressive pediatric malignancies, characterized by poor prognosis and limited therapeutic options. Emerging evidence suggests that specific lipid mediators significantly influence the growth, survival, and progression of this high-risk neuroblastoma subtype.

PGE2 was upregulated in liver by ARA (Table 2). It has been identified as a critical promoter of MYCN-amplified neuroblastoma growth and survival. PGE2 enhances MYCN-amplified neuroblastoma cell survival through EP4 receptor activation and subsequent cAMP/PKA signaling, which may stabilize MYCN protein and enhance its oncogenic functions (17). Inhibition of cyclooxygenase-2 (COX-2), the enzyme responsible for PGE2 synthesis, has shown promising anti-tumor effects in preclinical models of MYCN-amplified neuroblastoma, suggesting a potential therapeutic approach targeting this pathway (18). High dose DHA or EPA is predicted to have a similar effect on PGE2 production, blocking PGE2 production by outcompeting ARA for access to the COX-1 and COX-2 enzymes, while maintaining their function.

**Table 2.** ARA-derived eicosanoids in tissue. Ratio of DHA-dosed to ARA-dosed oxylipins for those that are p<0.05. Comparison of ARA and DHA dosed animal fold-changes compared to controls as described in the Methods. MRM transitions are reported as (precursor/product) masses. – means not significant The sole dihomo-gamma linolenic acid (20:3n-6)- derived eicosanoid, PGE1, is presented as the last entry. – means not significant

| ARA Derived Eicosanoids | Liver | Spleen | Brain | Skeletal Muscle |
| --- | --- | --- | --- | --- |
| PGE2 (351/271) | 0.18 | - | - | - |
| PGD2 (351/189) | 0.03 | - | - | - |
| PGF2a (353/193) | 0.06 | - | - | 0.33 |
| 8-iso-PGF2a (353/193) | - | - | - | 0.32 |
| TXB2 (369/169) | 0.04 | - | - | 0.34 |
| 5-HETE (319/115) | 0.09 | 0.32 | - | - |
| 12-HETE (319/179) | 0.12 | - | - | 0.38 |
| 15-HETE (319/219) | 0.14 | 0.32 | - | 0.34 |
| 20-HETE (319/289) | 0.11 | 0.12 | - | 0.16 |
| LTB4 (335/195) | - | - | - | - |
| LXA4/15(R)-LXA4 (351/115) | 0.14 | - | - | - |
| LXB4 (351/221) | - | - | - | - |
| 8,9-EET (319/127) | 0.10 | 0.21 | - | 0.35 |
| 11,12-EET | 0.08 | 0.30 | - | 0.37 |
| 14,15-EET | 0.07 | 0.35 | - | - |
| 5,6 DiHETrE (337/145) | 0.13 | 0.21 | - | 0.38 |
| 8,9 DiHETrE (337/127) | 0.24 | 0.20 | 0.20 | 0.14 |
| 11,12-DiHET | 0.25 | 0.16 | 0.33 | 0.23 |
| 14,15-DiHET | 0.28 | 0.19 | 0.32 | 0.27 |
| PGE1 (353/ 317) | 0.20 | - | - | - |

In contrast, 15-HETE demonstrates context-dependent effects in MYCN-amplified neuroblastoma, with several studies indicating potential anti-tumor properties. 15-HETE reportedly can induce growth arrest and apoptosis in MYCN-amplified neuroblastoma cell lines through activation of PPARγ signaling, which antagonizes MYCN-mediated transcriptional programs (19). This suggests a potential therapeutic avenue through modulation of 15-lipoxygenase activity or direct administration of 15-HETE derivatives. Our data show 15-HETE is significantly upregulated by ARA in liver, spleen, and skeletal muscle.

DHA-dosing upregulated the EPA-derived mediator 17,18-EpETE in all four tissues, and its hydrolysis product 17,18-DiHETE in liver and skeletal muscle, compared to ARA-dosing (Table 1). The 17,18-EpETE has demonstrated significant anti-tumor effects in MYCN-amplified neuroblastoma models. This epoxide can downregulate MYCN expression and inhibit neuroblastoma cell proliferation through modulation of PI3K/Akt signaling pathways, which are critical for MYCN protein stability. Similarly, its metabolite 17,18-DiHETE retains some anti-tumor activity but with reduced potency, implicating epoxide hydrolase activity in determining the efficacy of these mediators in neuroblastoma contexts (20).

Among the DHA-derived mediators (Table 3), 14-HDHA was upregulated in spleen, brain and skeletal muscle. It and its downstream metabolites have shown promising anti-tumor effects in MYCN-amplified neuroblastoma. 14-HDHA can inhibit neuroblastoma growth through multiple mechanisms, including reduction of inflammation in the tumor microenvironment and direct induction of apoptosis in neuroblastoma cells expressing high MYCN levels (21). The potential role of 14-HDHA as a precursor for maresin biosynthesis may contribute to its anti-tumor effects, though this remains to be fully elucidated.

**Table 3.** DHA-derived docosanoids in tissue. Ratio of DHA-dosed to ARA-dosed oxylipins for those that are p<0.05. Comparison of ARA and DHA dosed animal fold-changes compared to controls as described in the Methods. MRM transitions are reported as (precursor/product) masses. – means not significant

| DHA Derived Docosanoids | Liver | Spleen | Brain | Skeletal Muscle |
| --- | --- | --- | --- | --- |
| 7,8-EpDPA (343/113) | - | - | - | - |
| 10,11-EpDPA (343/153) | - | 2.37 | 1.33 | 1.83 |
| 13,14-EpDPA (343/161) | - | 2.77 | 1.53 | 2.61 |
| 16,17-EpDPA (343/273) | - | 1.97 | - | 2.27 |
| 19,20-EpDPA (343/241) | - | - | - | - |
| 7,8-DiHDPA (361/127) | - | - | - | - |
| 10,11-DiHDPA (361/153) | - | - | 1.86 | 2.12 |
| 13,14-DiHDPA (361/193) | - | 2.40 | 1.61 | 2.09 |
| 16,17-DiHDPA (361/233) | - | - | - | - |
| 19,20-DiHDPA (361/229) | - | 4.85 | - | 2.26 |
| 4-HDHA (343/101) | - | - | - | - |
| 7-HDHA (343/141) | - | - | - | - |
| 8-HDHA (343/125) | - | - | - | - |
| 10-HDHA (343/181) | - | 2.34 | 1.35 | 1.81 |
| 11-HDHA (343/165) | - | - | - | 1.69 |
| 13-HDHA (343/193) | - | 2.11 | - | 2.19 |
| 14-HDHA (343/205) | - | 2.85 | 1.64 | 2.61 |
| 16-HDHA (343/233) | - | - | - | - |
| 17-HDHA (343/245) | - | 1.91 | - | 2.31 |
| 20-HDHA (343/241) | - | - | - | - |

Thromboxane A2 (measured as TXB2) was upregulated by ARA in liver (Table 2). It has been implicated in promoting aggressive behavior in MYCN-amplified neuroblastoma. Elevated thromboxane production in MYCN-amplified tumors compared to non-amplified counterparts, with thromboxane receptor antagonists demonstrating efficacy in reducing tumor growth and metastasis in preclinical models (22). This suggests that targeting thromboxane signaling may represent a viable strategy for *MYCN*-amplified neuroblastoma treatment.

8,9-EET, 11,12-EET, and 14,15-EET were all upregulated by ARA in liver and spleen, and the former two in skeletal muscle (Table 2). The EETs, particularly 11,12-EET and 14,15-EET, appear to promote MYCN-amplified neuroblastoma growth through enhancement of angiogenesis and direct effects on tumor cell survival. Inhibition of soluble epoxide hydrolase, which increases EET levels, accelerated neuroblastoma growth in *MYCN*-amplified models, while EET antagonists reduced tumor burden (23). This indicates a complex role for EETs in neuroblastoma progression that may depend on MYCN status and tumor microenvironment factors.

### MYCN Non-Amplified Neuroblastoma

MYCN non-amplified neuroblastoma generally presents with better prognosis compared to its amplified counterpart, though high-risk subtypes still exist within this group. The influence of lipid mediators on this neuroblastoma variant demonstrates distinct patterns compared to MYCN-amplified disease.

PGE2 appears to exert less pronounced effects on MYCN non-amplified neuroblastoma compared to *MYCN*-amplified disease. While PGE2 can still promote cell survival and proliferation in non-amplified neuroblastoma cells, the magnitude of these effects is significantly reduced, suggesting that PGE2-mediated signaling may be partially dependent on MYCN amplification status (24). This differential sensitivity may provide therapeutic opportunities for stratified approaches targeting prostaglandin signaling based on *MYCN* status.

PGD2 and its downstream metabolites demonstrate significant anti-tumor properties in *MYCN* non-amplified neuroblastoma. PGD2 can induce apoptosis and cell cycle arrest in non-amplified neuroblastoma cells through activation of the DP1 receptor and subsequent induction of p53-dependent pathways (25). This effect appears more pronounced in *MYCN* non-amplified cells compared to *MYCN*-amplified counterparts, suggesting a potential therapeutic avenue for this specific neuroblastoma subtype.

DHA upregulated the EPA-derived mediator 18-HEPE in all tissues (Table 1). 18-HEPE and its downstream metabolites have shown promising anti-tumor effects in MYCN non-amplified neuroblastoma models. 18-HEPE inhibits tumor growth and enhances the efficacy of conventional chemotherapeutic agents in non-amplified neuroblastoma through modulation of inflammatory signaling in the tumor microenvironment. The potential role of 18-HEPE as a precursor for E-series resolvins may contribute to these effects, though direct evidence in neuroblastoma contexts remains limited.

20-HETE was upregulated in liver, spleen, and skeletal muscle (Table 2). It demonstrates pro-tumorigenic effects in MYCN non-amplified neuroblastoma through stimulation of angiogenesis and direct enhancement of tumor cell survival. Elevated 20-HETE levels in non-amplified neuroblastoma tissues compared to normal adrenal tissue, with inhibition of 20-HETE synthesis significantly reducing tumor growth in preclinical models (26). This suggests that targeting cytochrome P450-mediated 20-HETE production may represent a viable therapeutic strategy specifically for MYCN non-amplified disease.

Among the lipoxins, LXA4 demonstrates significant anti-tumor properties in *MYCN* non-amplified neuroblastoma. Paradoxically, it was upregulated in liver by ARA (Table 2). LXA4 can inhibit tumor growth and reduce metastatic potential in non-amplified neuroblastoma models through modulation of inflammatory signaling and direct effects on tumor cell migration (27). These effects appear to be independent of *MYCN* status, suggesting potential broad applicability across neuroblastoma subtypes.

### MYCN-Driven Cancers (Beyond Neuroblastoma)

MYCN amplification or overexpression occurs in various malignancies beyond neuroblastoma, including medulloblastoma, small cell lung cancer, and certain subtypes of neuroendocrine tumors. The influence of lipid mediators on these MYCN-driven cancers reveals important insights into potential therapeutic strategies.

ARA-derived PGE2 has been consistently identified as a promoter of MYCN-driven cancer progression across multiple tumor types. In MYCN-amplified medulloblastoma, PGE2 enhances tumor cell survival and resistance to conventional therapies through EP4 receptor-mediated activation of PI3K/Akt signaling, which stabilizes MYCN protein (28). Similar mechanisms have been observed in MYCN-driven small cell lung cancer, suggesting a conserved role for PGE2 in supporting MYCN oncogenic functions across different cancer contexts.

The ARA metabolite 5-HETE appears to support MYCN-driven cancer growth through promotion of angiogenesis and direct effects on tumor cell survival. Elevated 5-HETE levels in MYCN-amplified medulloblastoma compared to non-amplified tumors, with inhibition of 5-lipoxygenase significantly reducing tumor growth and extending survival in preclinical models (29). This suggests that targeting 5-HETE production may represent a viable strategy for MYCN-driven cancers beyond neuroblastoma.

Among the EPA-derived mediators, 17,18-EpETE and its metabolites have shown promising anti-tumor effects across multiple MYCN-driven cancer models. 17,18-EpETE can downregulate MYCN expression and inhibit tumor growth in MYCN-driven small cell lung cancer by modulating MAPK signaling pathways (30). Similar effects were observed in *MYCN*-amplified medulloblastoma, suggesting a conserved mechanism that may be exploited therapeutically.

The DHA-derived mediator 14-HDHA demonstrates significant anti-tumor properties in various *MYCN*- driven cancers. 14-HDHA can inhibit tumor growth and enhance the efficacy of conventional therapies in MYCN-amplified medulloblastoma by reducing inflammation in the tumor microenvironment and directly inducing apoptosis in tumor cells (21). These effects appear to be partially dependent on MYCN status, suggesting specificity for MYCN-driven cancer biology.

ARA-derived EETs, particularly 11,12-EET and 14,15-EET, have been implicated in promoting the growth of MYCN-driven cancers through enhancement of angiogenesis and direct effects on tumor cell survival. demonstrated that EET antagonists significantly reduced tumor burden in MYCN-amplified medulloblastoma models, suggesting a conserved role for these epoxyeicosanoids in supporting MYCN-driven cancer progression across different tumor types (23).

### MYC-Associated Cancers

*MYC* (*c-MYC*) dysregulation occurs in approximately 70% of human cancers, including Burkitt lymphoma, breast cancer, and multiple myeloma. The influence of lipid mediators on MYC-associated cancers reveals both similarities and differences compared to MYCN-driven malignancies.

ARA-derived PGE2 has been identified as a critical promoter of MYC-associated cancer growth and progression. In *MYC*-driven lymphomas, PGE2 enhances tumor cell survival and proliferation through EP4 receptor-mediated activation of cAMP/PKA signaling, which stabilizes MYC protein and enhances its transcriptional activity (31). Similar mechanisms have been observed in MYC-overexpressing breast cancer, suggesting a conserved role for PGE2 in supporting *MYC* oncogenic functions across different cancer contexts.

In contrast to its effects on *MYCN*-amplified cancers, ARA-derived 12-HETE demonstrates particularly pronounced pro-tumorigenic effects in MYC-associated malignancies. Elevated 12-HETE production in MYC-driven lymphomas compared to normal lymphoid tissue, with inhibition of 12-lipoxygenase significantly reducing tumor growth in preclinical models (32). Mechanistically, 12-HETE appears to enhance MYC protein stability through modulation of GSK-3β activity, representing a potential therapeutic target specific for MYC-associated cancers.

Among the EPA-derived mediators, 5-HEPE has demonstrated significant anti-tumor effects in MYC-associated breast cancer models. Research by revealed that 5-HEPE can downregulate MYC expression and inhibit tumor cell proliferation by antagonizing 5-HETE-mediated signaling pathways, which are critical for *MYC* protein stability in certain cancer contexts (33). This suggests a potential therapeutic avenue through dietary EPA supplementation or direct 5-HEPE administration in *MYC*-driven cancers.

The DHA-derived mediators 17-HDHA and 4-HDHA demonstrate significant anti-tumor properties in various MYC-associated cancers. These compounds can inhibit tumor growth in *MYC*-driven lymphoma models by reducing inflammation in the tumor microenvironment and directly inducing apoptosis in tumor cells (34). The potential roles of these mediators as precursors for D-series resolvins and maresins may contribute to their anti-tumor effects, though direct evidence in *MYC*-associated cancer contexts remains limited. iso-PGF2α, a marker of oxidative stress, appears to have complex and context-dependent effects in MYC-associated cancers. Elevated levels of this isoprostane have been observed in MYC-driven tumors, potentially reflecting the increased oxidative stress associated with MYC overexpression. While 8-iso-PGF2α itself may not directly influence tumor growth, it serves as an important biomarker for monitoring oxidative stress-targeted interventions in MYC-associated malignancies (35).

Lipoxins LXA4 and LXB4 demonstrate significant anti-tumor properties in MYC-associated cancers through modulation of inflammatory signaling and direct effects on tumor cell survival. Research by revealed that LXA4 can inhibit tumor growth and enhance the efficacy of conventional therapies in MYC-driven lymphoma through FPR2/ALX receptor activation and subsequent inhibition of NF-κB signaling, which is critical for MYC-mediated transcriptional programs (36). This suggests potential therapeutic applications for lipoxins or their stable analogs in MYC-associated malignancies.

### Lipid Mediator Summary

The contrasting roles of ARA-versus DHA-derived lipid mediators have emerged as a unifying theme in MYC/MYCN-driven oncogenesis. ARA metabolites such as PGE2, EETs, and 20-HETE consistently act in a pro-tumor capacity – they stimulate angiogenesis, proliferation, invasion, and immune evasion while blunting apoptosis. MYC-overexpressing cancers leverage these mediators to sustain their unchecked growth. In parallel, MYC and MYCN actively suppress the production of omega-3-derived lipid signals (for instance, by downregulating DHA synthesis in MYCN-amplified cells) as a means of disarming the cell’s intrinsic tumor-suppressive circuitry.

By contrast, mediators derived from DHA and EPA, including epoxide derivatives like 17,18-EEQ and 19,20-EDP, hydroxylated intermediates like 14-HDHA, and resolvins/maresins, exert broad anti-tumor effects. These omega-3 lipids oppose angiogenesis, re-sensitize cancer cells to apoptosis, and help normalize the inflammatory microenvironment to favor tumor control. Together, the evidence paints a cohesive biochemical picture: MYC-driven tumors thrive on a lipid signaling landscape dominated by ARA-derived “go” signals, whereas enhancing DHA-derived “stop” signals can tilt the balance toward growth arrest and regression.

This dualism suggests that therapeutic strategies combining MYC pathway inhibition with modulation of lipid mediators – for example, COX/CYP pathway blockers or pro-resolving lipid analogs – may hold particular promise in high-MYC cancers by simultaneously starving the tumor of pro-growth eicosanoids and reactivating dormant anti-tumor pathways. In sum, lipid mediators emerge not just as downstream metabolic bystanders but as active orchestrators of the MYC oncogenic program, with ARA- and DHA-derived signals serving as yin and yang in the tumor ecosystem.

## Conclusion

From a translational perspective, our results suggest dietary modulation of PUFA intake—specifically, reducing ARA and increasing DHA—may offer a complementary, low-toxicity approach to conventional neuroblastoma therapies. The substantial tumor suppression observed with DHA treatment suggests that omega-3 supplementation might be valuable in neuroblastoma management. This is particularly promising given that high-dose omega-3 supplementation is well-tolerated in humans for extended periods with only mild side effects (10).

Study limitations include the model’s incomplete representation of the human neuroblastoma’s complex tumor microenvironment, the use of fixed doses rather than titration, and the lack of detailed analysis of tumor-infiltrating immune cells and lipid mediator profiles. Future research should include comprehensive lipidomic profiling, detailed immunophenotyping, exploration of potential synergies with standard therapies, investigation of ROS-independent mechanisms, and evaluation in patient-derived xenograft models.

Within these limitations, our work demonstrates that arachidonic acid promotes, while docosahexaenoic acid suppresses, tumor progression in a MYCN-driven neuroblastoma model. Daily oral administration of ARA (75 µL/day; 4.6 g/day hEq) significantly accelerated tumor growth, while DHA (200 µL/day; 24 g/day hEq) substantially inhibited tumor development compared to control treatment. These findings align with our recent findings, where we demonstrated that ultra-high-dose omega-3 supplementation (EPA, DHA, or oxidation-resistant deuterated DHA) at human-equivalent doses of 12-14 g/day completely blocked tumor formation in a similar model. These results highlight the critical influence of dietary fatty acid composition on neuroblastoma behavior and underscore the therapeutic potential of omega-3 supplementation in neuroblastoma management. Given that ultra-high-dose omega-3 therapy has been successfully used and well-tolerated in humans for other conditions, our findings provide a compelling foundation for further investigation into dietary interventions as complementary approaches in pediatric oncology. Modulation of PUFA intake may represent a promising, low-toxicity strategy to influence disease outcomes in neuroblastoma patients, potentially allowing for reduced dosages of conventional genotoxic therapies and improved quality of life for survivors.

